# Probabilistic inference of bifurcations in single-cell data using a hierarchical mixture of factor analysers

**DOI:** 10.1101/076547

**Authors:** Kieran R. Campbell, Christopher Yau

**Affiliations:** Wellcome Trust Centre for Human Genetics University of Oxford

## Abstract

Modelling bifurcations in single-cell transcriptomics data has become an increasingly popular field of research. Several methods have been proposed to infer bifurcation structure from such data but all rely on heuristic non-probabilistic inference. Here we propose the first generative, fully probabilistic model for such inference based on a Bayesian hierarchical mixture of factor analysers. Our model exhibits competitive performance on large datasets despite implementing full MCMC sampling and its unique hierarchical prior structure enables automatic determination of genes driving the bifurcation process.

## 1 Introduction

Trajectory analysis of single-cell data has become a popular method that attempts to re-infer lost temporal information such as a cell’s differentiation state. Such analyses reconstruct a measure of a cell’s progression through some biological process, known as a *pseudotime*. Recently, attention has turned to modelling bifurcations where part-way along such trajectories cells undergo some fate decision and branch into two or more distinct cell types. Several methods have been proposed to infer bifurcation structure from single-cell data. Wishbone [1] constructs a *k*-nearest neighbour graph and uses shortest paths from a *root* cell to define pseudotimes, using inconsistencies over multiple paths to detect bifurcations. Diffusion Pseudotime (DPT) [2] similarly constructs a transition matrix where each entry may be interpreted as a diffusion distance between two cells. Bifurcations are inferred by identifying the anticorrelation structure of random walks from both a root cell and its maximally distant cell. While DPT arguably has a probabilistic interpretation, neither method specifies a fully generative model that incorporates measurement noise, while both infer bifurcations after constructing pseudotimes.

Here we propose a Bayesian hierarchical mixture of factor analysers for inferring bifurcations from single-cell data. Our model is unique compared to existing methods in the following: (1) by specifying a fully generative probabilistic model we incorporate measurement noise into inference and provide full uncertainty estimates for all parameters, (2) we simultaneously infer cell “pseudotimes” and branching structure as opposed to post-hoc branching inference as is typically performed, and (3) our hierarchical shrinkage prior structure automatically detects which features are involved in the bifurcation, providing statistical support for detecting which genes drive fate decisions. In the following, we introduce our model and apply it to both a toy dataset as well as previously published single-cell RNA-seq data. We finish with a discussion on the challenges of such models and future directions to be explored.

## 2 Model

We begin with an *N* × *G* matrix of (normalised) gene expression for *N* cells and *G* genes, where **y**_*i*_ denotes the *i*^*th*^ row vector corresponding to the gene expression for cell *i*. We assign a pseudotime *t*_*i*_ to each cell along with a binary variable *γ*_*i*_ indicating to which of *B* branches cell *i* belongs:

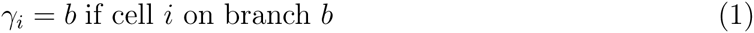

with *b* ∈ 1,…, *B*.

The pseudotime *t*_*i*_ is a surrogate measure of a cell’s progression along a trajectory while it is the behaviour of the genes - given by the factor loading matrix - that changes between the branches. We therefore introduce two factor loading matrices Λ_*γ*_ = [**c**_*γ*_ **k**_*γ*_]*, γ* 1*,…, B* for each branch modelled.

The likelihood of a given cell’s gene expression measurement conditional on all the parameters is then given by

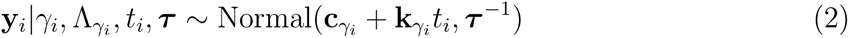

We motivate the prior structure as follows: if the bifurcation processes share some common elements then the behaviour of a non-negligible subset of the genes will be (near) identical across branches. It is therefore reasonable that the factor loading gradients **k***_γ_* should be similar to each other unless the data suggests otherwise. We therefore place a prior of the form

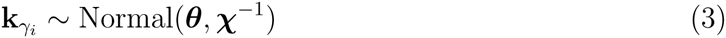

where ***θ*** denotes a common factor gradient across branches. This has similar elements to Automatic Relevance Determination (ARD) models with the difference that rather than shrinking regression coefficients to zero to induce sparsity we shrink factor loading gradients towards a common value to induce similar behaviour between mixture components. We can then inspect the posterior precision to identify genes involved in the bifurcation: if *χ*_*g*_ is very large then the model is sure that *k*_0*g*_ ≈ *k*_1*g*_ and gene *g* is not involved in the bifurcation; however, if *χ*_*g*_ is relatively small then |*k*_0*g*_ − *k*_1*g*_| ≫ 0 and the model indicates that *g* is involved in the bifurcation.

With these considerations the overall model becomes

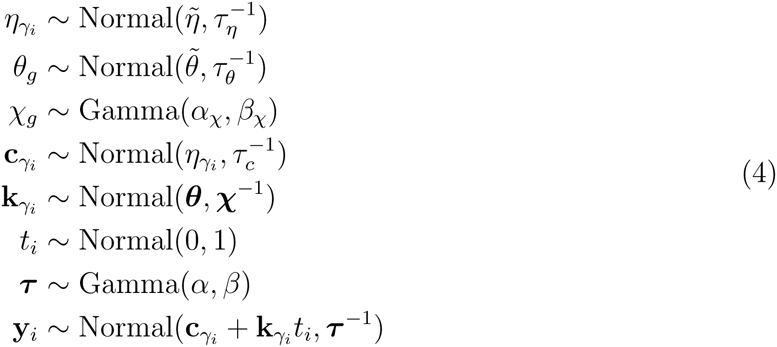

where 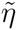, 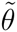, *τ*_*η*_, *τ*_*θ*_, *τ*_*c*_, *α*_*χ*_, *β*_*χ*_, *α* and *β* are hyperparameters fixed by the user. As the hierarchical model exhibits complete conjugacy inference is performed using Gibbs sampling.

## 3 Results

### 3.1 Toy dataset

We first demonstrate our method on a synthetic ‘toy’ dataset of 100 bifurcating cells and 40 genes, half of which exhibit differential behaviour across the bifurcation and half of which show similar behaviour. Our synthetically generated data is mildly mis-specified with respect to our model to demonstrate robustness when using real genomic data. For example, the generated gene behaviour across pseudotime is nonlinear, albeit monotonic and pseudotimes generated from Unif[0, 1).

The Gibbs sampling algorithm for this dataset takes less than one minute to run on a laptop computer for 2 × 10^5^ iterations. The results can be seen in figure 1. Figure 1A shows a principal components representation of the data in which the distinctive bifurcation pattern can be seen. Points are coloured by the maximum a-posteriori (MAP) estimates of the pseudotimes, which clearly shows a trajectory running parallel to principal component 1 (PC1) of the data. Figure 1B shows the model infers the MAP estimates of the branch indicator variables *γ*_*i*_ correctly, with uncertainty prior to the bifurcation point followed by the probability “collapsing” to 0 or 1 as the branches become apparent. Note that branch assignment prior to the bifurcation is essentially meaningless and as long as we observe the correct gene behaviour along pseudotime across branches the actual assignment is of no real concern.

**Figure 1:**
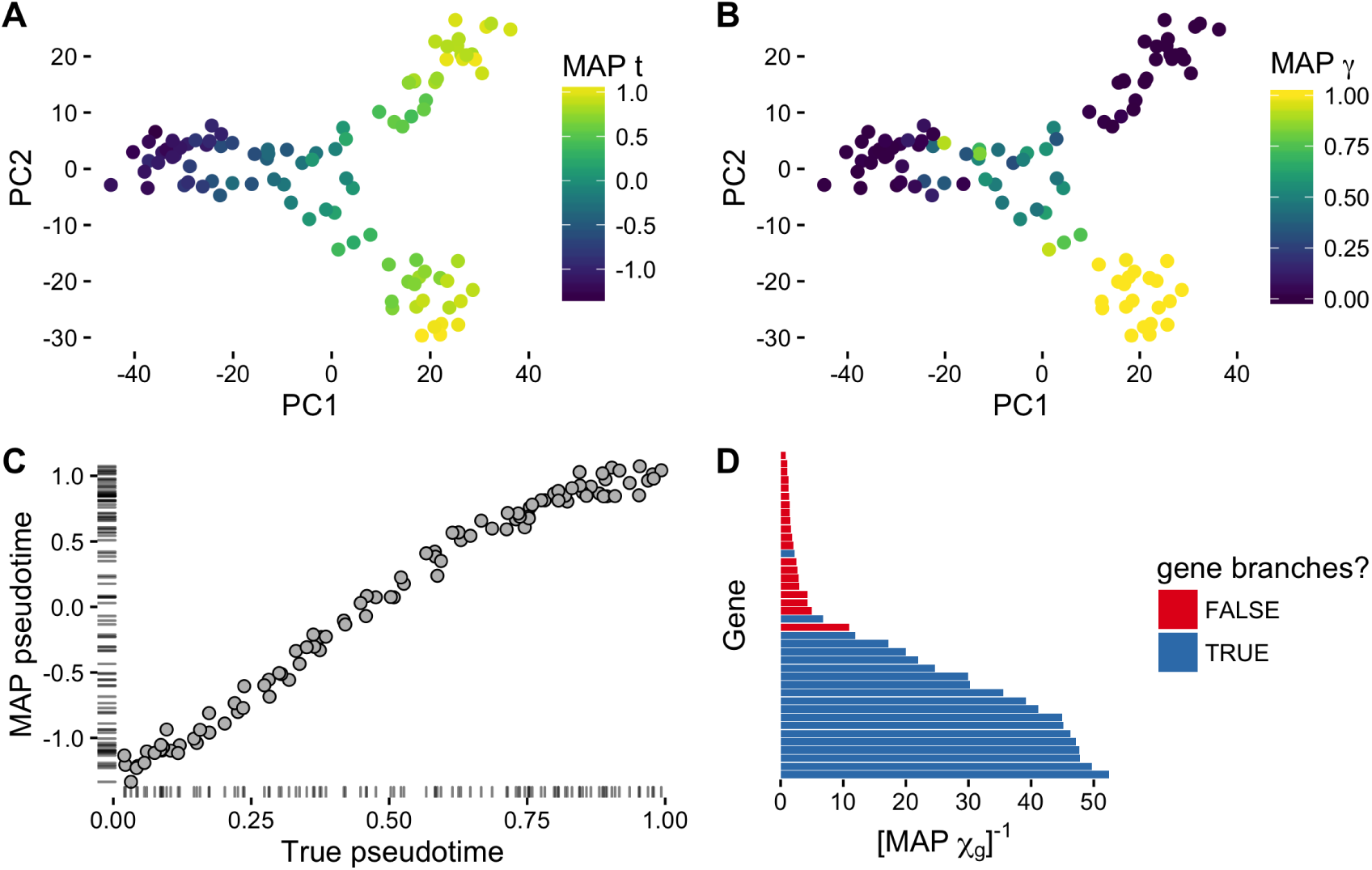
Probabilistic inference of bifurcations in toy data.

We subsequently compared the inferred pseudotimes to the true (generated) ones. Figure 1C shows good agreement, with a (pearson/spearman) correlation *ρ* > 0.98. The slight warping of the relationship towards the end of pseudotime may be a consequence of generating data with a nonlinear likelihood yet performing inference using a linear model. Finally, figure 1D shows the inverse of the MAP estimates of *χ*_*g*_, an indicator of a gene’s involvement in the bifurcation process. Of the genes used in the toy dataset, half showed differential behaviour across the bifurcation and half shows common behaviour. The MAP estimates of *χ*_*g*_ clearly segregate the two types of genes, suggesting this method may perform well at determining which genes drive cell fate processes.

### 3.2 Single-cell RNA-seq data

We next applied our method to previously published single-cell RNA-seq data of HSPCs differentiating into myeloid and erythroid precursors [3]. To reduce the dataset to a computationally feasible size we selected the most highly variable genes passing a pre-set threshold, resulting in 69 retained genes across 4423 cells. Inference using Gibbs sampling took around 15 minutes on a laptop computer for 5 × 10^4^ iterations. The results can be seen in figure 2(A-B). The MAP pseudotime estimates clearly recapitulates the trajectory in the data as shown using a tSNE representation from [1], while the MAP estimates of *γ*_*i*_ detects the branching structure in the data, consistent with previous methods.

**Figure 2:**
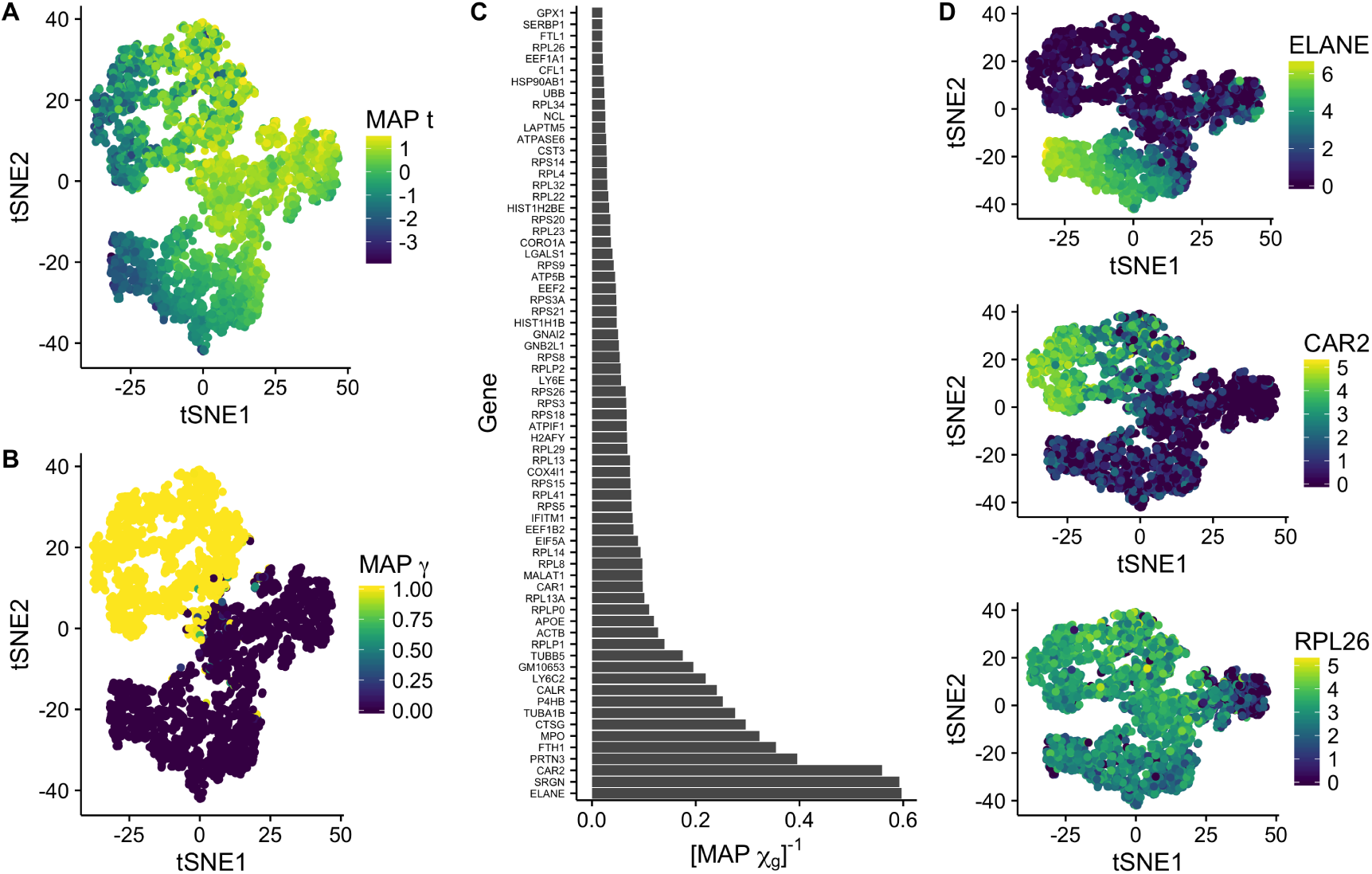
Inference of bifurcations in scRNA-seq data.

We went on to analyse the genes suggested by the model to be involved in the bifurcation process. Figure 2C shows the inverse posterior mean of *χ*_*g*_, with larger values indicating more evidence that gene *g* is involved in the bifurcation process. For illustration purposes, we plot the expression of *ELANE* and *CAR2*, which the model suggests will show differential behaviour across the bifurcation, along with *RPL26*, which the model suggests will show common behaviour (figure 2D).

## 4 Discussion

In this paper we have presented a Bayesian hierarchical mixture of factor analysers for inference of bifurcating trajectories in single-cell data. While the model performs well on both real and synthetic data, inference is sensitive to parameter initialisation, akin to selecting a *root* cell in previous methods. Indeed, it is easy to reason that there are three maxima of the posterior space that are indistinguishable without the addition of further information.

There are several extensions that can be applied to our model. For example, a zero-inflated likelihood, as in [4], could be considered to reduce the model mis-specification. Furthermore, while the model performs well on even the largest of current single cell RNA-seq datasets, inference via Variational Bayes may need to be considered in the future. This would be practical given the model’s conjugate exponential family structure.

## Acknowledgments

K.C. is supported by a UK Medical Research Council funded doctoral studentship. C.Y. is supported by a UK Medical Research Council New Investigator Research Grant (Ref. No. MR/L001411/1), the Wellcome Trust Core Award Grant Number 090532/Z/09/Z, the John Fell Oxford University Press (OUP) Research Fund and the Li Ka Shing Foundation via a Oxford-Stanford Big Data in Human Health Seed Grant.

